# GWAS identifies candidate genes controlling adventitious rooting in *Populus trichocarpa*

**DOI:** 10.1101/2022.06.14.496209

**Authors:** Michael F. Nagle, Jialin Yuan, Damanpreet Kaur, Cathleen Ma, Ekaterina Peremyslova, Yuan Jiang, Christopher J. Willig, Greg S. Goralogia, Alexa Niño de Rivera, Megan McEldowney, Amanda Goddard, Anna Magnuson, Wellington Muchero, Li Fuxin, Steven H. Strauss

## Abstract

Adventitious rooting is critical to the propagation, breeding, and genetic engineering or editing of trees. The capacity for plants to undergo these processes is highly heritable; however, the basis of its genetic variation is largely uncharacterized. To identify genetic regulators of these processes, we performed a genome-wide association study (GWAS) using 1,148 genotypes of *Populus trichocarpa*. GWAS are often limited by the abilities of researchers to collect precise phenotype data on a high-throughput scale; to help overcome this limitation, we developed a computer vision system to measure an array of traits related to adventitious root development in poplar, including temporal measures of lateral and basal root length and area. GWAS was performed using multiple methods and significance thresholds to handle non-normal phenotype statistics, and to gain statistical power. These analyses yielded a total of 277 unique associations, suggesting that genes that control rooting include regulators of hormone signaling, cell division and structure, and reactive oxygen species signaling. Genes related to other processes with known roles in root development, and numerous genes with uncharacterized functions and/or cryptic roles, were also identified. These candidates provide targets for functional analysis, including physiological and epistatic analyses, to better characterize the complex polygenic regulation of adventitious rooting.

## Introduction

The species within the genus *Populus spp.* (poplar) are among the most rapidly-growing trees of the northern hemisphere (Dickmann & Stuart, 1983) and have outsized roles in natural ecosystems as keystone species (Brunner *et al*., 2004; Kouki *et al*., 2004; Bailey & Whitham, 2006; Kivinen *et al*., 2020). They are also of major economic importance for agroforestry, and as sources of wood, fiber, and biofuel (Sun *et al*., 2021; Fuertes *et al*., 2021). In either context, the growth and asexual propagation of poplar relies on the rapid establishment, proliferation and maintenance of a robust root system for nutrient and water absorption. Elite hybrid clones of poplar are propagated in stool beds through a process that relies on the ability of cuttings to undergo adventitious rooting (Stanton *et al*., 2019). In addition, asexual reproduction of poplar in nature commonly occurs via the process of “root sprouting” from existing roots or the root-shoot junction zone (Wiehle *et al*., 2009). Moreover, biotic and abiotic stresses such as waterlogging and pest damage often interfere with above-ground tree health by damaging root systems (Brandt *et al*., 2003; Štícha *et al*., 2016), a stress that is certain to be exacerbated by climate change (Overpeck & Udall, 2020; Gullino *et al*., 2021). A deeper understanding of the genes that control rooting may provide new insights into means for improved propagation of recalcitrant genotypes and species, options for improvement of regeneration during genetic engineering/editing, and suggest new strategies for mitigating stress in managed and wild populations.

Adventitious rooting in poplar is a highly complex trait that is regulated by many factors, including plant age, genotype, and physiology; the many forms of plant stress; and environmental cues such as temperature, photoperiod, and nutrients. These act through phytohormone signaling cascades that lead to differentiation and development of root tissue. Overexpression and RNAi-mediated suppression of over two dozen genes involved in phytohormone synthesis and response have been reported to lead to increased or decreased adventitious root formation and/or root growth (reviewed by Bannoud & Bellini, 2021). These root-related traits have been shown to be genotype-dependent, with phenotypic variation across *Populus spp.* depending in large part on variable sequence and expression of phytohormone-related genes, especially those involved in auxin pathways or crosstalk between auxin and other phytohormones (Ribeiro *et al*., 2016; Sun *et al*., 2019). Genome-wide association studies (GWAS) provide opportunity for insight into how variation in root-related traits across genotypes results from variation in these phytohormone regulators and other genes. To date, GWAS of adventitious rooting performed in plants including rice (Ribeiro *et al*., 2016) and *Populus* (Sun *et al*., 2019) have contributed to an improved understanding of these gene-function relationships and others.

GWAS of root traits, both from adventitious roots and non-adventitious roots, commonly involve measurement of the lengths, diameters, or types of roots, among other statistics. As collection of these traits can prove laborious and time-consuming, a wide array of computer vision tools have been developed to extract these features from root images, with varying degrees of human intervention or automation (e.g., Arsenault *et al*., 1995; Das *et al*., 2015; Zhang *et al*., 2018; Yasrab *et al*., 2019; Zheng *et al*., 2020). These methods have been applied to enable GWAS and other genetic analyses of rooting in common bean, cowpea (Burridge *et al*., 2016), maize (Arsenault *et al*., 1995; Zhang *et al*., 2018) and hybrid poplar (Sun *et al*., 2019), among others. Methods involving manual measurements with rulers or ImageJ have been applied to GWAS of rooting traits in plants including maize (Trachsel *et al*., 2011), rice (Courtois *et al*., 2013; Li *et al*., 2017; Wang *et al*., 2018; Xu *et al*., 2020; Zhang *et al*., 2020), wheat (Ayalew *et al*., 2018) and hybrid poplar (Dash *et al*., 2018). The production of more general and user-friendly root phenotyping platforms can help to extend automated methods to more species and laboratories, potentially assisting expansion of GWAS population size by reducing the amount of manual labor needed for phenotyping. With increased population size, GWAS gain greater statistical power and improved ability to detect significant effects of genes regulating traits, particularly those or smaller effect size, and those imparted by rare alleles (López-Cortegano & Caballero, 2019).

Here, we report insights into the genetic control of rooting obtained through GWAS. By use of a large resequenced and highly polymorphic population of wild *Populus trichocarpa* that shows very rapid decay of linkage disequilibrium—and a novel machine vision phenomic system and multiple GWAS pipelines—we were able to statistically detect large numbers of candidate genes. The potential for rare allele discovery was enhanced by our use of the SNP-Set Sequence Kernel Association Test, which upweights and combines effects from rare SNPs to increase power in their detection (Wu *et al*., 2011). We report a total of 277 unique associations passing significance thresholds, including many involved in hormone signaling, cell division, and post-translational modification of proteins—in addition to many genes of unknown function.

## Methods and Materials

### Plant materials

We used a *P. trichocarpa* GWAS population that was recently expanded to include a total of 1,323 genotypes (Yates *et al*., 2021); subsets of this population were used in previous GWAS projects (Tuskan *et al*., 2018; Bdeir *et al*., 2019; Weighill *et al*., 2019; Chhetri *et al*., 2020; Chen *et al*., 2021; Nagle *et al*., 2022). This population is comprised of variation in wild *P. trichocarpa* spanning regions of British Columbia, Washington, Oregon, Idaho and northern California. Clone banks for this GWAS population were produced in multiple locations, among which a replicate in a Corvallis, OR field location was utilized to obtain cuttings for this study. This study was performed using materials collected as described in previous work (Nagle *et al*., 2022). In summary, dormant cuttings were collected in the winters of 2018, 2019 and 2020, then rooted up to a year later and grown in a greenhouse. Finally, fresh cuttings were collected and frozen for 2-4 weeks, then used for rooting assays.

### Assay of rooting

Cuttings were placed in 50mL Falcon tubes with water and allowed to root. Beginning two weeks later, images were collected at weekly timepoints for four weeks. Prior to taking each image, plants were removed from water and placed on top of a surface with roots arranged to separate putative lateral and basal roots. To aid their recognition by our machine vision pipeline, basal roots were laid downward atop blue felt while lateral roots were laid to the side on gray felt. Each image also included a label and ruler. Plants were imaged from above using a Canon Rebel XSi DSLR camera attached to a mount and facing downward. Due to practical limitations in the number of cuttings that could be studied at once, the study was divided into eight “phases,” each of which featured ∼400 cuttings, including two replicates for each of ∼200 given genotypes.

### Computer vision pipeline

We adopted the DeepLab network (Chen *et al*., 2018) with backbone ResNet50 (He *et al*., 2016) as our segmentation model for its efficiency and accuracy. We trained two different segmentation models: the first was used to segment an image into background, plant, ruler, and label (Model 1); the second was used to segment the image into background, leaf, stem, and root (Model 2). Below, we introduce how we collected training labels to train the two networks and then used the two networks to measure biological traits of interest.

Because images were collected in various orientations, with the camera in either portrait or landscape mode, we first rotated images to a uniform orientation. Next, the background was segmented based on color thresholding. As the plant, ruler and label varied in color and were found in approximately similar positions from image to image, we first segmented these components based on their spatial positioning and colors using mean-shift segmentation and k-means clustering. Images successfully segmented as such were used to produce a training set with approximately 500 images used to train a deep model for segmentation of the remaining images. Inference was performed using this model, and correct examples were used to retrain the model with an expanded dataset, resulting in a final model trained with 2,239 examples (Deep Model 1). Afterward, to segment the plant into stem, root and leaf, we performed mean-shift segmentation and applied a location threshold to produce a training set of approximately 900 images. These were used to train a second deep model and the training set was again expanded by running inference on new images and selecting correct results, resulting in a final training set with 3,496 training examples (Deep Model 2).

Following training of both deep segmentation models, they were applied for inference of the full dataset. All images were standardized to the same orientation, followed by deployment of both models. The final result of segmentation was separation of background, leaf, stem, root, ruler, and label for each image. Next, the segmentation results were further analyzed to produce statistics on biological traits of interest. Since the camera height varied across images, we computed the number of pixels per ruler width for each image to enable standardization via the actual size represented by a single pixel.

We proceeded to compute the lengths and area of roots in centimeters, as well as the diameters of stems. Root statistics were computed as follows. (1) First, we isolated the segment of root by distinct connections to the stem via connected components (2) The background was classified as top background and bottom background based on the color of the felt background below the stem, allowing each root to be classified as lateral or basal depending on the background. (3) For each basal and lateral root, we computed root length using a distance map and root area by counting pixels. Longest root length (LRL) and total root area were computed separately for basal and lateral roots in each image. RGB and false-color images of the segmentation process are shown in Fig. 1.

**Figure 1.**
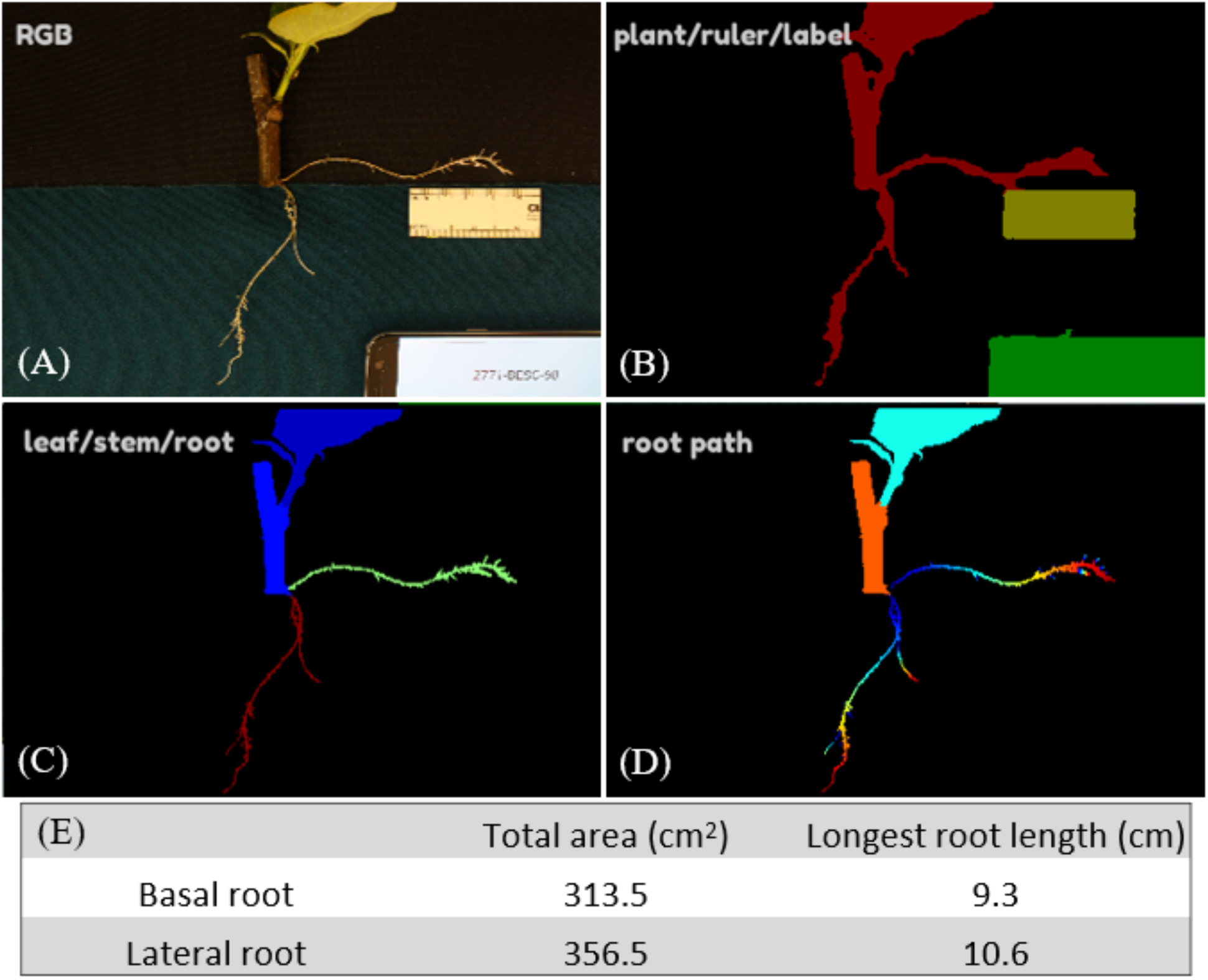
Workflow for phenotyping root traits: (A) RGB images were collected for plants with rulers and labels. (B) The first round of segmentation was performed to separate the plant, ruler and label. (C) A subsequent round of segmentation separated the roots by their type (basal or lateral, shown respectively in red and green). (D) Finally, the length of each root was measured from root tip to connection to stem. (E) An example of total area and longest root length computed over basal and lateral types of adventitious roots.

### Data Error Checking Methods

Source images were compared to segmented images (e.g. Fig. 1) to identify cases where data contained errors for the length, area, type or number of roots. Each image was scored according to the error or errors observed (Table 1). Images were then sorted into folders according to the types of data errors they contained using a spreadsheet and an R script.

**Table 1.**
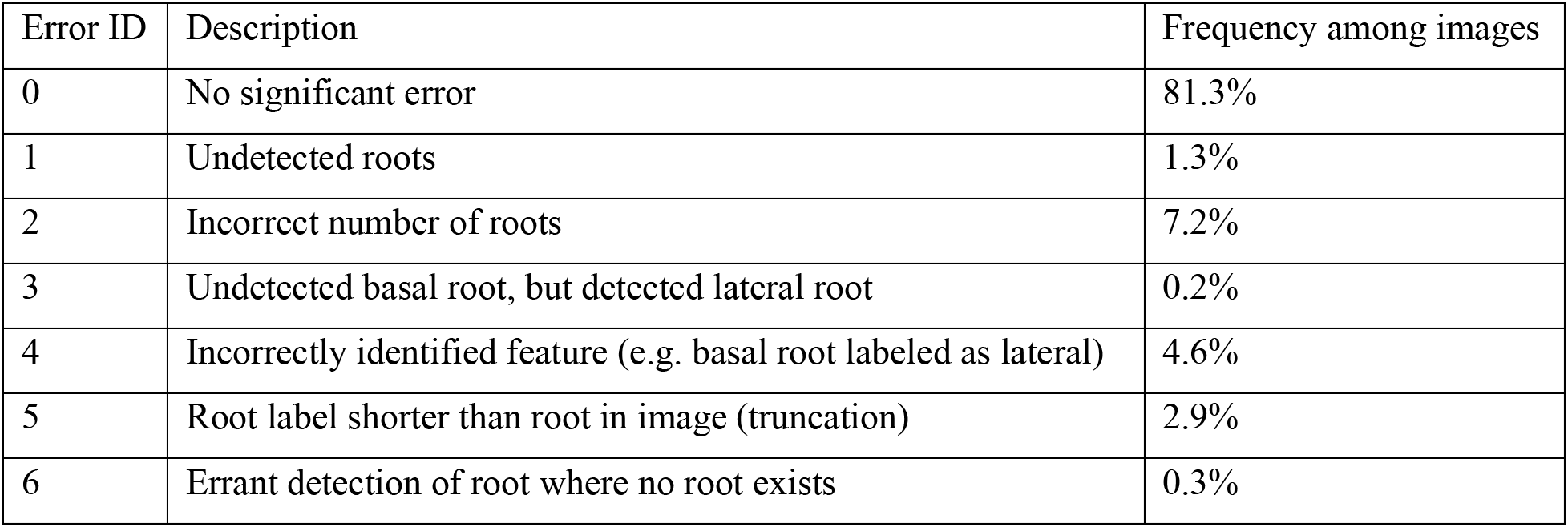
Descriptions of error types and their frequencies.

| Error ID | Description | Frequency among images |
| --- | --- | --- |
| 0 | No significant error | 81.3% |
| 1 | Undetected roots | 1.3% |
| 2 | Incorrect number of roots | 7.2% |
| 3 | Undetected basal root, but detected lateral root | 0.2% |
| 4 | Incorrectly identified feature (e.g. basal root labeled as lateral) | 4.6% |
| 5 | Root label shorter than root in image (truncation) | 2.9% |
| 6 | Errant detection of root where no root exists | 0.3% |

### Data Correction Methods

ImageJ version 1.53a software and the SmartRoot version 4.21 plugin were installed and used to make all measurements. Source images were opened in ImageJ with the SmartRoot plugin then converted to an 8-bit greyscale and inverted. The global scale was then set to 1 pixel/unit in ImageJ and 2.54 DPI in SmartRoot to acquire units of pixels for all measurements. Roots that required correction of length data were then traced using the SmartRoot Trace Root tool. To calculate root areas, the image color threshold was adjusted until only the roots were highlighted, then the area was selected using the Wand Tracing tool and measured in ImageJ. Root types were corrected based on manual judgement of whether a root was a lateral root or basal root, while recalling that these were placed on different areas of felt during imaging as previously described.

### Data preparation

Mean values of each trait were computed across replicates for each genotype and used for downstream GWAS. Principal component analysis (PCA) was performed using ‘stats::princomp’ in R to produce PCs representing trends across traits and timepoints. PCA was performed over three batches of traits: 1) area of basal or lateral roots at all timepoints; 2) LRL for basal or lateral roots at all timepoints; and 3) all traits, including area and LRL of basal or lateral root at all timepoints. Scree plots were consulted to determine the numbers of PCs, representing significant variation for each batch, to be used for downstream GWAS.

Normality of traits was assessed in parallel to traits for our published GWAS of *in planta* regeneration in poplar, using the same methods (Nagle *et al*., 2022). In short, we assessed normality of untransformed traits using Q-Q plots, histograms, Shapiro-Wilks tests and Pearson correlation coefficients with theoretical normal distributions, then applied necessary transformations including Box-Cox transformations, rank-based inverse normal transformations, removal of zero values and removal of outliers on a case-by-case basis for conservative but adequate transformation of each trait (Figure S2, Table S1-2).

### Association mapping

Association mapping of all traits was performed in parallel with traits in our published GWAS of *in planta* regeneration, using methods and SNP sets detailed in this previous work (Nagle *et al*., 2022). To summarize, four GWAS methods were used: 1) Genome-wide Efficient Mixed Model Association (GEMMA; Zhou & Stephens, 2012); 2) Generalized Mixed Model Association Test (GMMAT; Chen *et al*., 2016); 3) Fixed and Random Model Circulating Probability Unification (FarmCPU; Liu *et al*., 2016), specifically the implementation FarmCPUpp (Kusmec & Schnable, 2018); and 4) SNP-set (sequence) Kernel Association Test (SKAT; Ionita-Laza *et al*., 2013) with the Multi-Threaded Monte Carlo SKAT (MTMC-SKAT) R extension we developed. Versions of trait data transformed toward normal distributions were analyzed with GEMMA and FarmCPU, while binarized traits were analyzed with GMMAT and untransformed traits with MTMC-SKAT using resampling to avoid violations of linear model assumptions for high-confidence associations. GEMMA and GMMAT were run using kinship matrices to adjust for population stratification, with a set of 13.2 million SNPs with a minor allele frequency (MAF) threshold of 1% and that are missing in no more than 10% of genotypes. FarmCPU uses a novel approach to adjust for population stratification while avoiding overcorrection and was run with a SNP set with MAF above 5%, missing rate below 10% and pruning based on linkage disequilibrium (LD). MTMC-SKAT was run using six principal components derived from SNP data to correct for population stratification and a set of 34.0 M SNPs with missing rate below 15%. MTMC-SKAT was deployed on the high-performance cluster COMET, made available through NSF XSEDE (Towns *et al*., 2014). Stem diameter and phase were used as covariates for all GWAS methods.

To identify QTLs from results that are statistically significant, we computed multiple testing correction thresholds using the Bonferroni method (parameters: α = 0.05, *N* tests equal to *N* SNPs) and Benjamini-Hochberg false discovery rate (α = 0.10). We further sought to identify candidate genes that failed to meet significance according to either of these criteria, but were represented by a peak of QTLs showing a pattern of LD decay suggestive of a causative association. Toward this end, we applied an implementation of the Augmented Rank Truncation (ART; Vsevolozhskaya *et al*., 2019) over GEMMA and GMMAT results as we previously described (Nagle *et al*., 2022).

## Results

### Principal components describe complex patterns of root development

Significant patterns of root development over time and across root types (basal and lateral) were summarized by PCA. For each of the three batches of traits used for PCA, the top two PCs appear to explain significant portions of variance as indicated by Scree plots. PCA was performed over three batches as previously described (Methods and Materials). The first two batches, for root area or root length traits, each across basal and lateral roots and timepoints, both produced a first PC representing lateral root development independent of basal root and a second PC representing a lack of basal root development independent of lateral root (Fig. 2, Fig. S1). For the third batch, including all area and length traits together, the first PC shows a trend of overall root development and the second shows a preference for development of basal root rather than lateral root (Fig. S2).

**Figure 2.**
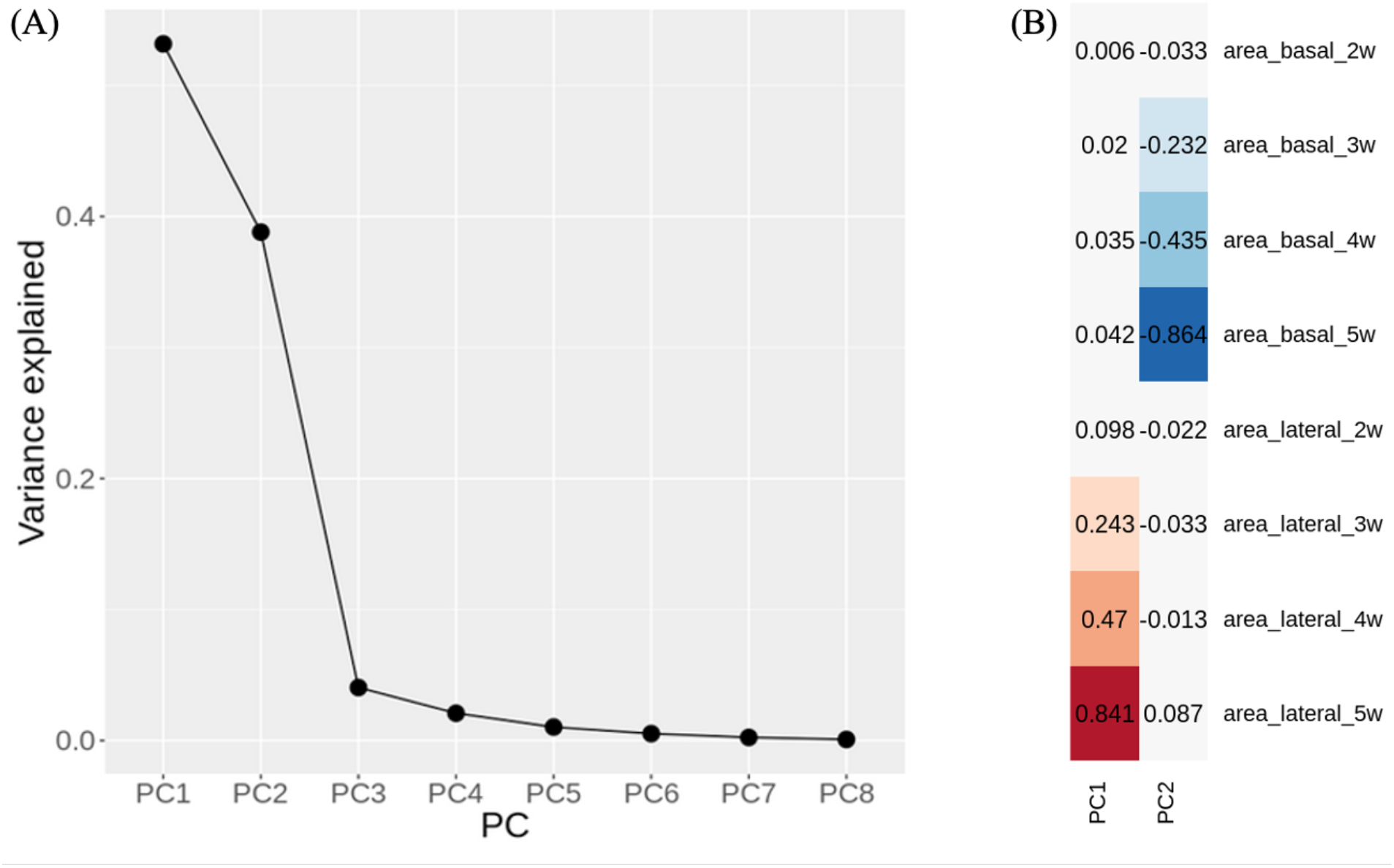
Results from PCA over all root area traits, across root type (basal or lateral) and all four timepoints of data collection: (A) Scree plot showing proportion of variance explained by each PC; (B) Loadings for top two PCs.

### Use of multiple GWAS methods yielded numerous associations

We evaluated results to identify candidate QTLs and genes implicated for all 13 traits with h^2^SNP above 0.10, three of which were evaluated with two different transformations due to ambiguity in the ideal transformation (Table S1-S3). We term a “QTL peak” as any SNP or SNP window associated with a given trait that is not within 30kb of any other SNP or SNP window with a lower *p*-value for the associated trait, thus appearing as the peak position of a group of signals on a Manhattan plot. Only MTMC-SKAT yielded QTL peaks passing the conventional Bonferroni threshold under the assumption that each test (SNP window) is independent. MTMC-SKAT using 3kb SNP windows yielded a total of 31 unique Bonferroni-significant associations across all traits, 22 of which had a window center within 5kb of an annotated gene, as well as 164 unique associations passing the FDR (α = 0.10) and/or the conservative Bonferroni threshold (112 of which had window centers within 5kb of a gene). The apparent power of SKAT is in contrast to GEMMA and GMMAT; GEMMA yielded only two associations passing the FDR (α = 0.10) threshold and none passing the conservative Bonferroni threshold, while GMMAT yielded none passing either. Statistical power for GEMMA and GMMAT was greatly increased by the use of ART to combine signals across windows of SNPs, which enabled detection of over 100 associations for GEMMA and eight for GMMAT (Fig. 3-4). Fig.5 provides an example of manual inspection of QTL peaks using integrative genomics viewer (IGV; Thorvaldsdóttir *et al*., 2013). Associations of note that are addressed in the discussion section are presented in Table 2, with summary statistics for all associations in Table S4 and details presented in Table S5-8. No significant associations were found using FarmCPU.

**Figure 3.**
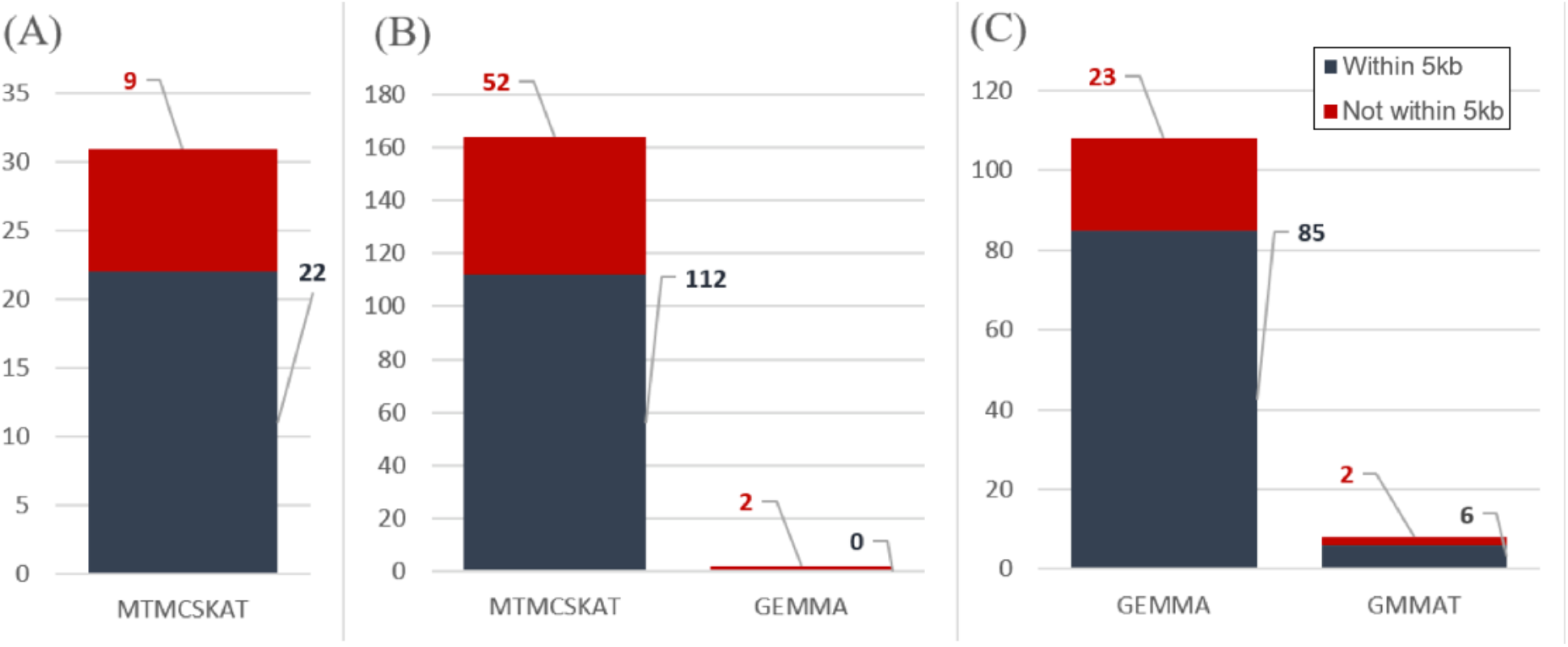
Barplots summarizing the numbers of associations from each GWAS method, with each type of significance threshold, as well as within a 5kb distance threshold to the nearest gene. QTL peaks were taken as the point with the lowest *p*-value at any given position near a significant SNP, where multiple points within the same peak may otherwise pass a given significance threshold. (A) QTL peaks passing the conservative Bonferroni threshold, given an assumption of independence of all SNP associations; (B) QTL peaks passing Benjamini-Hochberg threshold (FDR; α = 0.10) and/or the conservative Bonferroni Threshold. (C) QTL peaks passing ART-Bonferroni threshold (α = 0.05, N of # 1kb windows in genome); ART was only applied to the single-SNP GWAS methods, GEMMA and GMMAT.

**Figure 4.**
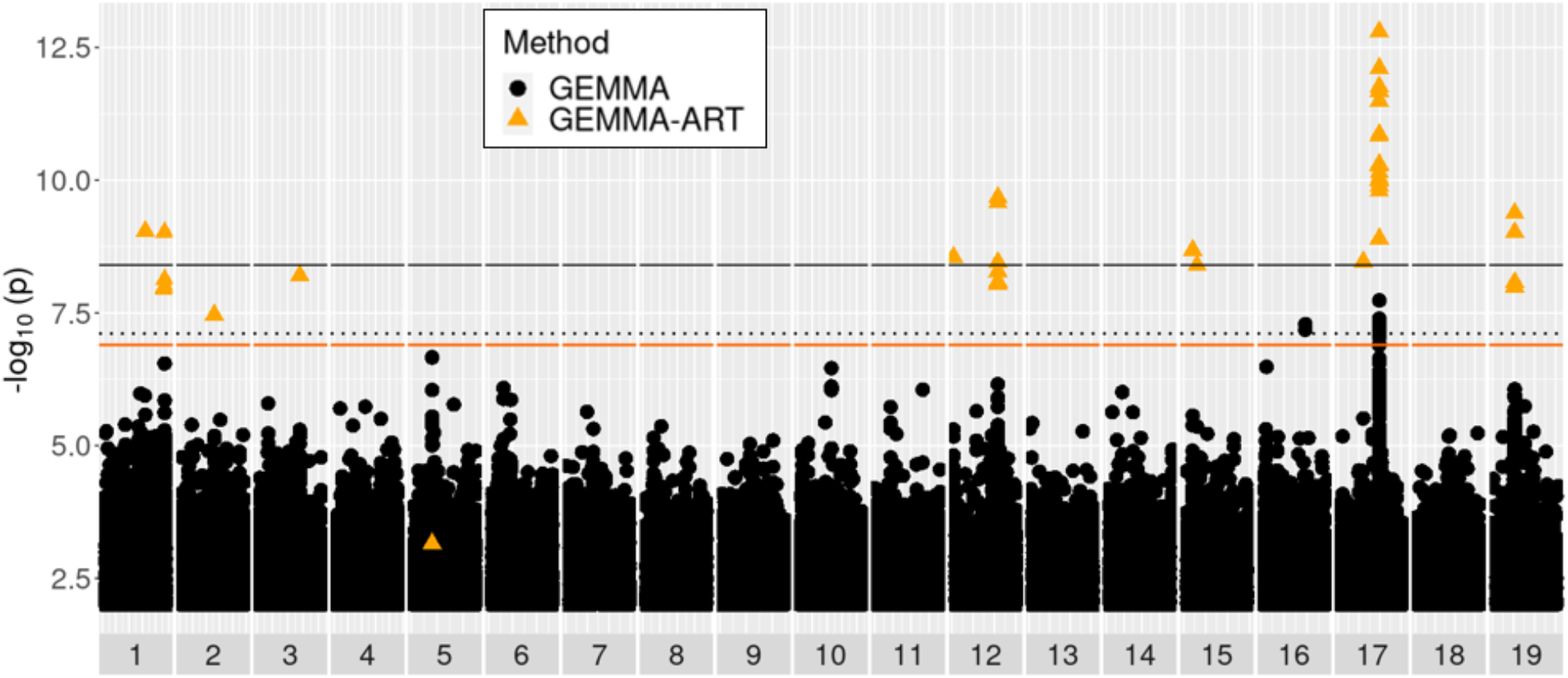
Manhattan plot of GEMMA results for the trait of longest lateral root length at week 3. Black and orange solid lines represent Bonferroni significance thresholds for GEMMA results with independent SNPs, and for ART applied to GEMMA over 1kb windows of SNPs. The black dotted line represents the significance threshold with a false discovery rate of 10% for independent SNP tests. Black circles represent tests of individual SNPs by GEMMA. Orange triangles represent 1kb windows tested by ART applied to GEMMA results.

**Figure 5.**
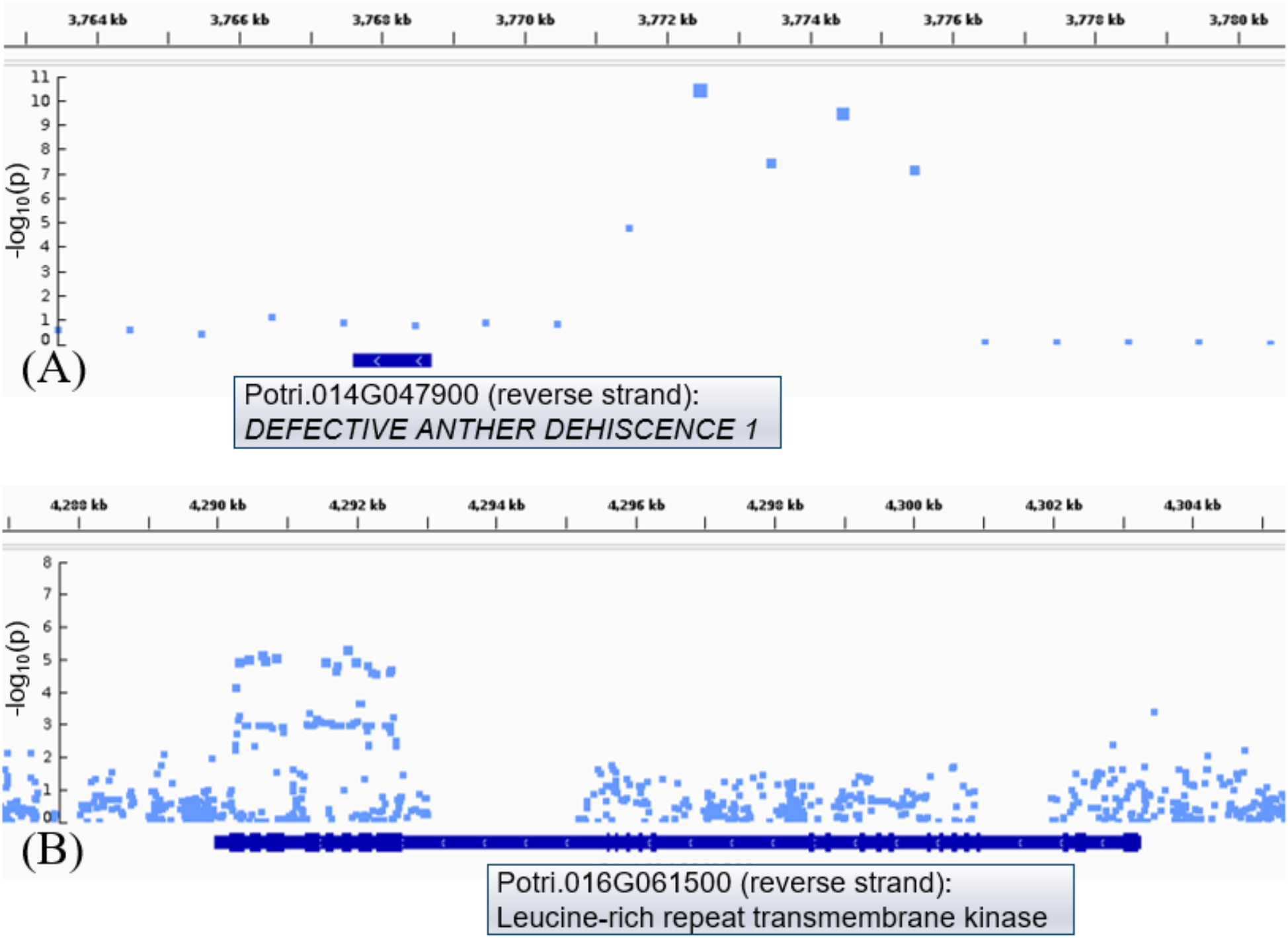
Close-up view of Manhattan plots aligned with *P. trichocarpa* genome annotation (v3.1) using IGV. (A) SKAT results (prior to resampling top associations with MTMC-SKAT) for LRL at week three display an association with a possible promoter region of a putative homolog of *DEFECTIVE ANTHER DEHISCENCE 1;* (B) GEMMA results (without GEMMA-ART displayed) for basal root area at week five display an association with a ∼2kb region of exons and short introns of a putative leucine repeat rich transmembrane protein kinase. Exons are visualized as thickened portions on the gene track. Plots are displayed without MTMC-SKAT and GEMMA-ART *p*-values for simplicity; these statistics can be found in Table S6-7. Boxes with gene accession IDs, strand of gene and gene info were added manually to IGV plots.

**Table 2.**
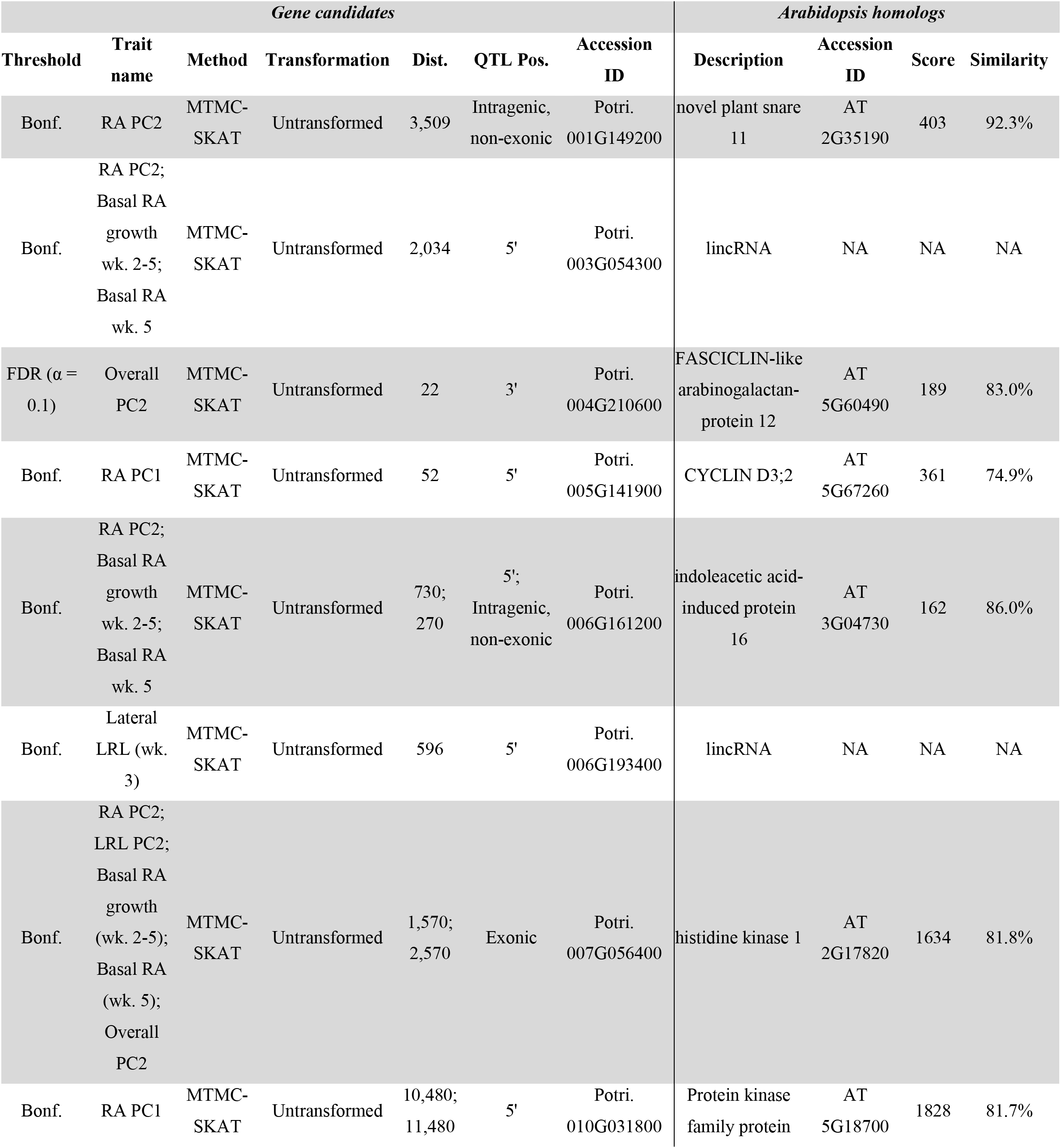
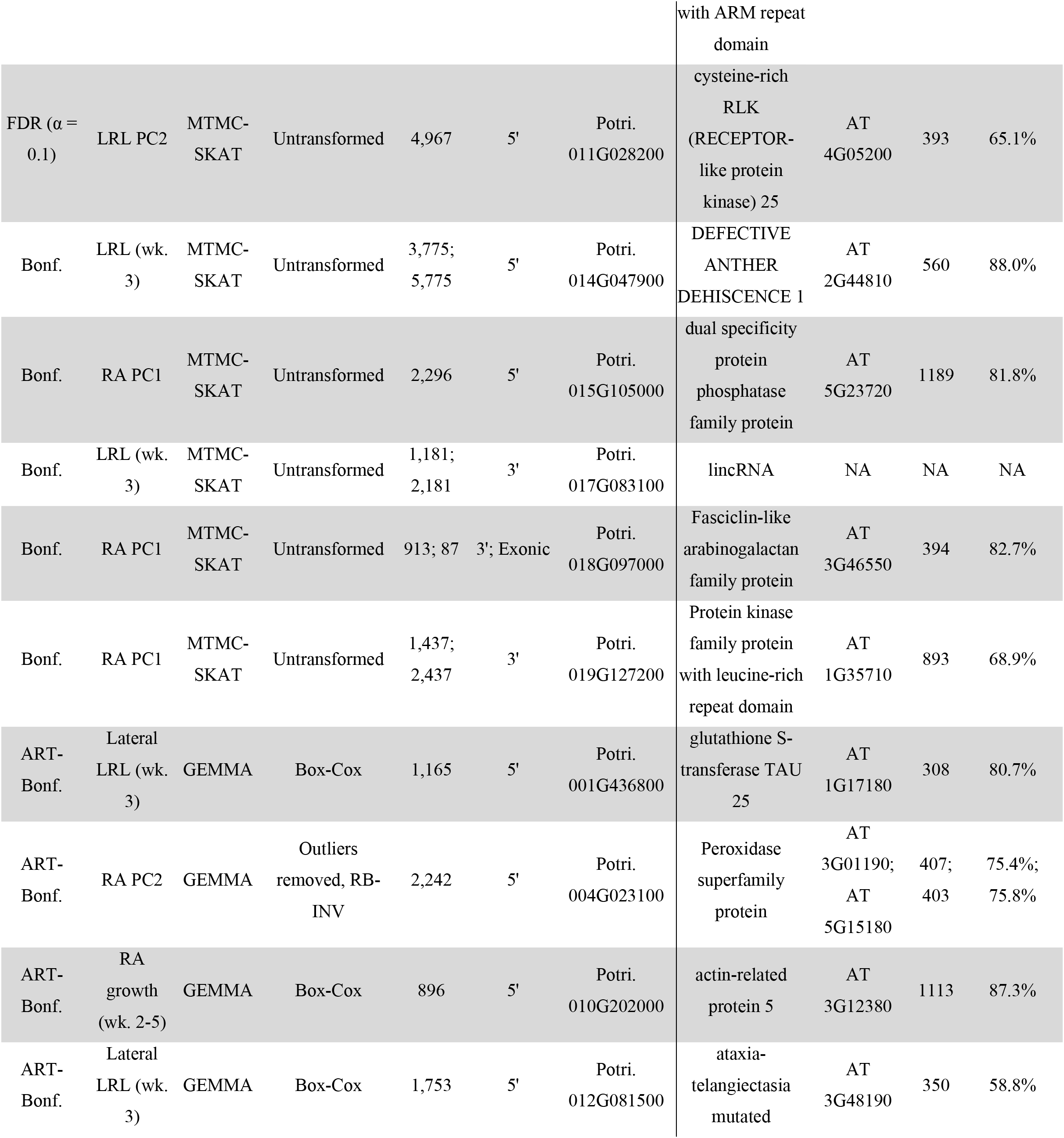
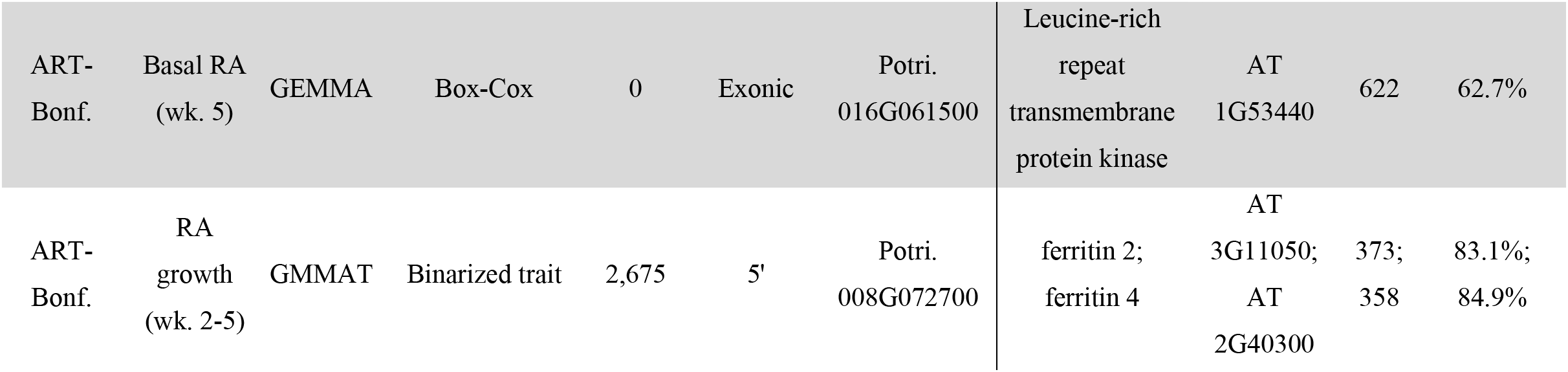
Twenty gene candidates with Arabidopsis homologs or encoding lincRNAs with putative role in biological processes of root development. For each candidate with an Arabidopsis homolog, relevant literature is summarized (Discussion). QTLs were identified from various traits related to root area (RA) and longest root length (LRL), for either basal or lateral ARs, or both types combined. For some associations found by MTMC-SKAT, two nearby SNP windows have equal *p*-values for association (1*10^-7^) due to the practical limits of our permutation analysis. Remaining associations are presented in Tables S5-S8.

## Discussion

As discussed in depth below, we reported a high number of putative associations from our GWAS pipeline. FarmCPU, however, gave no significant associations, and only a single candidate association in our prior study using this pipeline (Nagle *et al*., 2022). We are unaware of any other reports of this GWAS method being used in poplar. We speculate that the lack of success with this method may stem from the high SNP density and rapid linkage disequilibrium (LD) decay in poplar, the unique approach FarmCPU employs in controlling for population stratification using a limited set of SNPs, and/or the LD-based pruning that was needed to avoid errors during this process for the SNP set we used.

### Two putative modes of adventitious root regeneration in poplar

We observed that when ARs grew from the base of cuttings, the base was often enlarged, distorted, and disorganized in appearance, resembling the calli found in *in vitro* tissue cultures or at *in planta* wound sites (Fig. 6). We term these “basal ARs.” Although we are aware of little research into this specific type of AR in poplar, research in *Populus balsamifera* (a closely related species interfertile with *P. trichocarpa*), suggested that AR development depends on calcium and pH, and that hard callus may inhibit root emergence (Cormack, 1965). More recently, basal ARs growing from callus have been studied in *Pinus* (Rasmussen *et al*., 2009). These contrast with ARs growing from the sides of cuttings (“lateral ARs”), which appeared to grow directly from the stem without an intermediate callus stage (Fig. 6). Prior histological research in poplar indicated that lateral ARs appear to originate from secondary meristematic tissue in the cambium (Rigal *et al*., 2012). We therefore set up our phenomics pipeline to measure lateral and basal ARs separately, assuming they are biologically distinct and likely originate from different progenitor cells. Their high degree of independence was supported by our PCA analysis, which by inspection of loadings appear to represent “ratios” between the two types of root (Fig. 2, Fig. S1-2). There was also a number of GWAS associations for one root type or the other, or for these PCs (discussed below).

**Figure 6.**
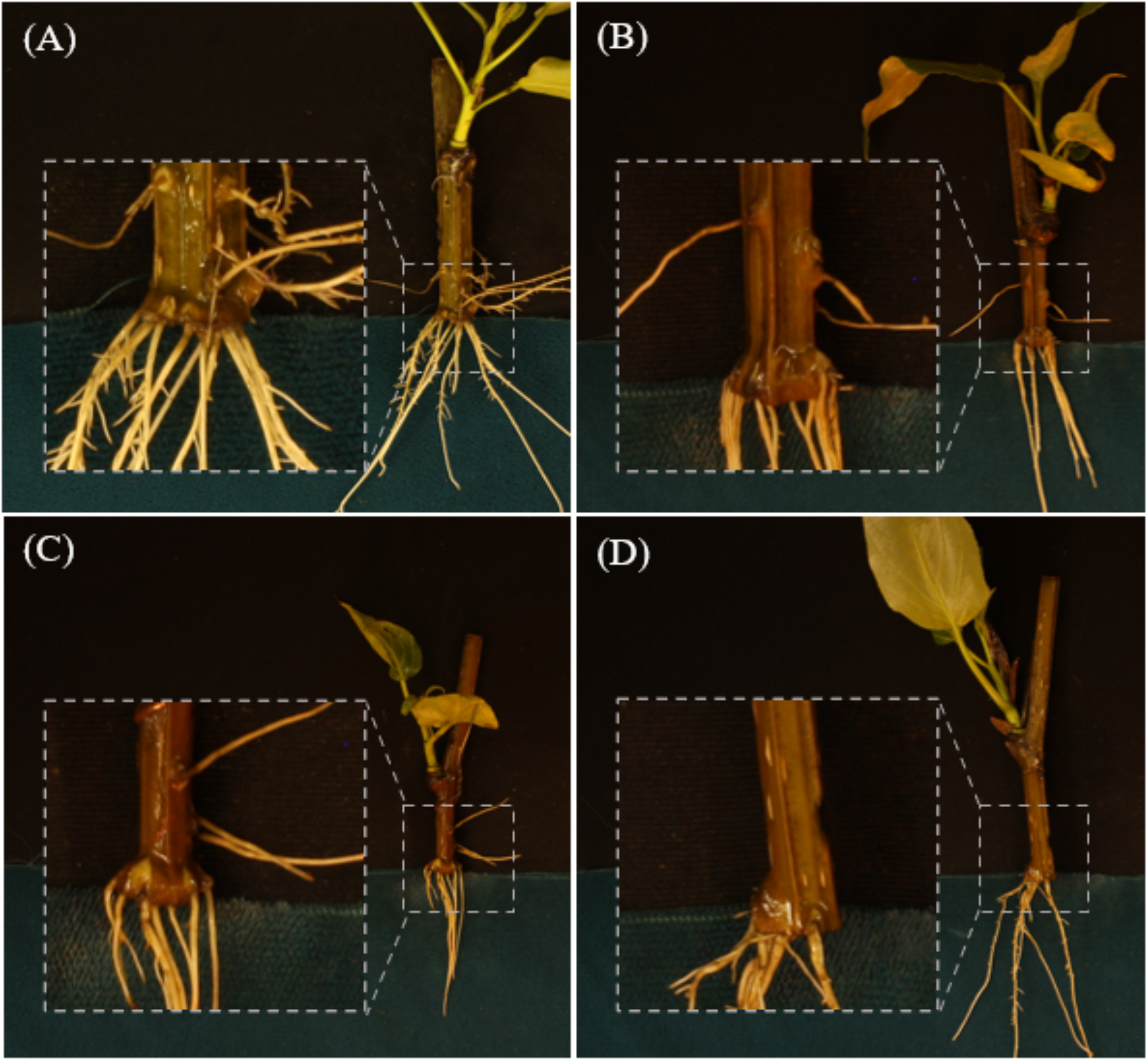
Selected images collected during root phenotyping, with zoomed-in views showing callus or callus-like tissue at the base of stems from which basal ARs emerge. (A) Genotype BESC-153, (B) BESC-327, (C) GW-9914, and (D) BESC-337.

### Phenomics workflow accelerated phenotyping but required human intervention

Our phenomics workflow using machine vision enabled us to extract root trait data from a number of images that would otherwise have been infeasible for humans unassisted. The training of models for this workflow did not require manual preparation of ground-truth semantic labels by humans using an annotation interface, but rather utilized ground-truth labels prepared by thresholding and cropping. Although this approach offers the advantage of accelerated training dataset preparation, it lacks the ability for machine vision results to be compared to human-produced ground truth labels using statistics such as intersection over union (IoU). To assess model performance, we inspected each image and grouped them according to notable errors that we observed. The most frequent errors were in root counting, found in at least 7.2.% of images (Table 1). Root count statistics were not used in GWAS because of the relatively high error rate and because the summary statistics of aggregate root area provides a proxy for root system proliferation, while being more robust to errors. Another common type of error was of incorrectly applied labels, for example basal roots labeled as lateral roots or vice versa (Table 1). With our system, we attempted to facilitate the labeling of basal and lateral roots by manually placing roots of each type in areas with different background colors (Fig. 6). We are not aware of other root phenomics workflows that aim to distinguish basal and lateral ARs, although several distinguished root type for non-adventitious roots, such as for alfalfa (Xu *et al*., 2022), Arabidopsis, wheat and *Brassica* (Yasrab *et al*., 2019).

We attempted to correct errors using ImageJ with the SmartRoot plugin (Methods and Materials). This error correction was performed on approximately 18.7% of images. Considering our hybrid approach utilizing both machine vision and manual correction of errant labels, our method is comparable to RootReader (Clark *et al*., 2013), a tool that similarly performs initial labeling of images and then allows for user correction. The scope of our work did not include the development of a user-friendly and generalizable root annotation method that can be practically applied by other labs with diverse imaging conditions and across diverse species of plants. However, several such tools have been recently developed, including RootNav 2.0, which has demonstrated an ability via transfer learning to generalize across different background types and species including maize, Arabidopsis and *Brassica* (Yasrab *et al*., 2019).

### Gene candidates represent diverse functional roles

#### Regulators of cell division and structure

D-type cyclins regulate the G1-to-S progression of the cell cycle and comprise a family of ten known proteins in Arabidopsis and 22 in poplar (Dong *et al*., 2011). Potri.005G141900 encodes a homolog of CYCLIN D3;2 (CYCD3;2) or CYCLIN D3;3 (CYCD3;3), which are among at least five D-type cyclins known to have roles in root precursor cells in embryonic and/or mature tissues (De Veylder *et al*., 1999; Nieuwland *et al*., 2009; Collins *et al*., 2012; Forzani *et al*., 2014). A comparative transcriptomic study in *Populus* found this gene to be differentially expressed between two genotypes with contrasting rates of stem growth and biomass accumulation (Han *et al*., 2020). In Arabidopsis, concurrent knockout of all three members of the CYCD3 clade led to retarded seed development, while overexpression of CYCD3;1 led to premature, irregular and disorganized division of the hypophysis (Collins *et al*., 2012). Transcription of CYCD3;1 and CYCD3;3 is negatively regulated by WUSCHEL-ASSOCIATED HOMEOBOX 5 (Forzani *et al*., 2014), a poplar homolog of which enhances AR in poplar when overexpressed (Li *et al*., 2018). Although we are unaware of any reports of mutant phenotypes resulting from CYCD3 overexpression or knockout in mature roots, several relevant root-related phenotypes have been reported for CYCD2 and CYCD4 members. CYCLIN D4;1 knockout was reported to reduce pericycle cell divisions as well as the number of lateral roots, while these phenotypes were rescued by exogenous auxin, perhaps due to auxin-responsiveness of D-type cyclins with overlapping functional roles (Nieuwland *et al*., 2009). CYCD2;1 overexpression led to increased root apical meristem divisions and increased sensitivity to effects of exogenous auxin in promoting lateral root formation, while knockout led to reduced auxin sensitivity although no reduction in RAM divisions, possibly due to redundant homologs (Sanz *et al*., 2011).

Potri.001G149200 encodes a homolog of NOVEL PLANT SNARE 11 (NPSN11), which is believed to interact with other SNARE proteins to provide energy needed for the fusion of membranes that give rise to the cell plate during cytokinesis (Zheng *et al*., 2002). Knockout of NPSN11 alone yields no mutant phenotype in Arabidopsis, putatively due to redundancy with a similar SNARE protein. Defects in cytokinesis and embryo development are conferred by simultaneous knockout of NSPN11 and the functionally redundant SNAP33 (El Kasmi *et al*., 2013).

Potri.004G210600 and Potri.018G097000 encode members of the FASICLIN-LIKE ARABINOGALACTAN (FLA) family, which consists of 21 members in Arabidopsis and has established roles in adhesion in cell walls, plasma membranes and extracellular matrices (reviewed by Zang *et al*., 2015). Transcriptomic analysis of tension wood development in *Populus* provides support for a role of Potri.004G210600 in cell wall structure (Bygdell *et al*., 2017). Knockout of Arabidopsis FLA4, also known as SALT OVERSENSITIVE 5, was reported to reduce root elongation, cell wall thickness and root tip swelling under salt stress (Shi *et al*., 2003) and experiments with ethylene inhibitors suggest that FLA4 functions downstream of ethylene signaling (Xu *et al*., 2008). We are unaware of reports of root-related mutant phenotypes of other FLA members.

Cell expansion is influenced by bundling and rearrangements of actin filaments, regulated at least in part by auxin and cytokinin (Zhu & Geisler, 2015; Scheuring *et al*., 2016; Arieti & Staiger, 2020). We report two gene candidates encoding putative actins or actin-like proteins, Potri.010G202000 and Potri.012G081500. In Arabidopsis, loss-of-function mutants of ACTIN 7 displayed reduced root length and reduced cell divisions in the proximal meristem (PM) of root tips as well as an increased number of transition zone (TZ) cells, while dual mutants of ACTIN 2 and ACTIN 8 presented a loss of root hairs and increases in TZ cells with a lack of PM effects (Kandasamy *et al*., 2009; Takatsuka *et al*., 2018). The variable effects of different auxins on root development, together with the lack of clear homology for these actin-related gene candidates, welcome physiological investigation of the mechanisms by which these gene candidates may affect root in poplar.

#### Regulators of hormone signaling

Potri.006G161200 encodes a member of the Aux/IAA F-Box protein family, believed to have 35 members in *Populus trichocarpa* (Kalluri *et al*., 2007) and 29 in Arabidopsis (Overvoorde *et al*., 2005), and appears to be a homolog of IAA16 or another member of the 29-gene IAA family. In Arabidopsis, Aux/IAA proteins have been well-characterized and are known to undergo auxin-dependent proteasomal degradation and to function via auxin-dependent protein-protein interactions that repress the transcriptional activity of various Auxin Response Factor (ARF) family members (reviewed by (Luo *et al*., 2018)). A specific role for ARF members in regulating AR development is believed to function via the action of downstream *GRETCHEN HAGEN* family genes responsible for conjugating jasmonic acid (JA) into bioactive jasmonoyl-L-isoleucine (JA-Ile). Evidence for conservation of this pathway in *Populus* has been reported, with enhanced or delayed AR development respectively resulting from overexpression or knockdown of a homolog of TRANSPORT INHIBITOR RESPONSE 1, responsible for Aux/IAA proteasomal degradation and found to interact with a *Populus* homolog of IAA28 (Shu, et al. 2019).

Further evidence for a role of JA signaling in rooting of poplar is indicated by an association with Potri.014G047900, encoding a homolog of DEFECTIVE IN ANTHER DEHISCHENCE1 (DAD1), which catalyzes the first step of JA biosynthesis (Ishiguro *et al*., 2001). Several possible roles for JA in adventitious rooting of *Populus* have been discussed in a recent review; in summary, these roles may include cross-talk with auxin signaling among others, and are evidenced to vary across genera (Bannoud & Bellini, 2021). Our previous GWAS of *in planta* regeneration in poplar support a major role for JA signaling in callus and shoot regeneration (Nagle *et al*., 2022; and sources cited within), via pathways that are likely to be relevant to adventitious rooting considering the previously discussed emergence of basal ARs from callus.

#### Regulators of post-translational modifications

Among our candidate genes, we report a notable number of genes encoding putative catalysts of post-translational modifications (PTMs), including a histidine kinase (Potri.007G056400), serine/threonine kinases (Potri.011G028200, Potri.010G031800, Potri.019G127200 and Potri.016G061500), a serine/threonine phosphatase (Potri.015G105000) and a glutathione-*S*-transferase (Potri.001G436800). We are particularly unsure of the precise mechanisms by which these candidates affect root traits because their PTM activity may be highly nonspecific and a majority of plant genes are likely to undergo PTMs. Evidence has been found for PTMs of over 12,000 substrates in Arabidopsis (Xue *et al*., 2022), including arabinogalactans (Schultz et al 2004) and microtubule proteins involved in cell structure and division (Parrotta *et al*., 2014) as well as hormone signal regulators (reviewed by Hill, 2015). Specific interactions between PTM catalysts and other gene candidates can be interrogated via statistical tests for epistasis.

Arabidopsis mutants of certain PTM-related candidate homologs display phenotypes relevant to root development. Mutants of the serine/threonine phosphatase PROPYZAMIND-HYPERSENSITIVE 1 (homolog of Potri.015G105000) demonstrate embryo fatality (for null mutants) or microtubule defects leading to left-handed helical growth of roots in seedlings (for mutants with reduced phosphatase activity) (Naoi and Hashimoto, 2004). HISTIDINE KINASE 1 (homolog of Potri.007G056400) is believed to have a role in abscisic acid (ABA) signaling, indicated by increased sensitivity of seedlings to the effects of ABA in inhibiting germination (Tran *et al*., 2007). In *Populus*, the putative histidine kinase Potri.007G056400 was found to be differentially expressed in *Populus* roots in response to boron deficiency (Su *et al*., 2019) and the putative serine/threonine kinase Potri.015G105000 was previously identified in GWAS as an association with bud set and growth period (McKown *et al*., 2014).

#### Noncoding RNAs

We sought to identify possible targets of putative ncRNAs that were found as associations in our GWAS, and indicated by the GreeNC pipeline to be probable ncRNAs (Di Marsico *et al*., 2022). Potri.003G054300 shares significant homology with the predicted exon of Potri.006G260300, a gene that appears to encode a transmembrane protein but for which we were unable to identify a homolog in model species. We did not find any known protein-coding genes that Potri.006G193400 and Potri.017G083100 align well with. We note that noncoding RNAs may be involved in processes other than RNA interference of coding genes, such as in ribonucleoprotein complexes and chromatin modification (reviewed by Statello *et al*., 2021). Whereas gene-silencing effects of ncRNAs can be predicted by sequence alignment, other roles may not be unraveled without wet-lab protocols (reviewed by Lucero *et al*., 2021).

#### Regulators of reactive oxygen species (ROS) signaling

ROS can affect or correspond to root development through multiple mechanisms. High levels of ROS are associated with various biotic and abiotic stressors in plants (Sharma *et al*., 2019; Qamer *et al*., 2021) and are well-established as a cause of DNA and tissue damage across eukaryotes (Arfin *et al*., 2021). As a means of post-transcriptional regulation, ROS can catalyze activation of deactivation of transcription factors (Wu *et al*., 2012; Kong *et al*., 2018) and other developmental genes such as cell cycle regulators (Yi *et al*., 2014). Additionally, the previously discussed roles of auxin in root signaling relate to ROS as the bioactive auxin IAA is produced by a peroxisome-mediated reaction involving the precursor IBA, producing nitric oxide as a byproduct (reviewed by Damodaran & Strader, 2019). Moreover, nitric oxide is involved in nitrosylation of proteins including the auxin receptor TRANSPORT INHIBITOR RESPONSE (TIR1), promoting its interaction with Aux/IAA proteins (Terrile *et al*., 2012) such as the previously discussed IAA16. NO and ROS also have roles in mediating symbioses with mycorrhizae as well as pathogen defense (reviewed by Martínez-Medina *et al*., 2019) although these roles are likely not relevant to our root assays in our laboratory using water rather than soil.

Oxidative stress is mitigated in part by ferritins, proteins that sequester Fe and thus prevent Fe from reacting with oxygen and producing oxygen radicals. Potri.008G072700 encodes a homolog of the four-member ferritin family in Arabidopsis, and is most closely related to *FERRITIN 2* (greater Smith-Waterman alignment score) and *FERRITIN 4* (greater residue similarity). The latter homolog has been studied in the context of root system architecture. While increasing concentrations of Fe in media led to an increase in lateral root density, this effect was abolished in triple knockouts of FERRITIN 1, 2 and 4 (Reyt *et al*., 2015).

In addition to catalyzing a PTM of endogenous proteins as previously discussed, glutathione *S-*transferases (GSTs) such as Potri.001G436800 are well-known to have roles in detoxification of xenobiotics such as herbicides, and have lesser-characterized roles in regulation of redox balance via glutathione, an antioxidant (reviewed by Hernández Estévez & Rodríguez Hernández, 2020). Transgenic studies have found Arabidopsis lines overexpressing various GST homologs to have increased tolerance to oxidative stress (Sharma *et al*., 2014; Xu *et al*., 2017) and enhanced root proliferation, although more research is needed to determine the specific mechanism or mechanisms by which GSTs regulate root development, whether via ROS, PTMs or other roles (Chen *et al*., 2012).

Other enzymes have more direct roles in oxidative stress and signaling, such as peroxidases that catalyze redox reactions (Yoshida *et al*., 2003). Potri.004G023100 is an example of a putative peroxidase and is closely related to Arabidopsis accessions AT3G01190 and AT5G15180, both of which are highly expressed in root apices (Klepikova *et al*., 2016) but have not been characterized in mutant studies to our knowledge.

### Agreement with previous GWAS of adventitious rooting in poplar

Very few of our gene candidates were also identified as possible regulators of adventitious root traits in published studies. In a prior GWAS in *Populus deltoides x simonii* that employed 434 genotypes and yield 224 QTLs, with multiple possible candidate genes being proposed for a given QTL, an uncharacterized gene believed to be a transcription factor (Potri.005G154200), as well as a putative xyloglucan endotransglucosylase/hydrolase (Potri.016G098600), were associated with rooting traits. They appeared as associations with lateral LRL at week three in our work, and with total number of roots in this prior work (Table S6). Potri.015G026500 encodes an uncharacterized putative phospholipase that is found as associated with lateral LRL at week 3 in our work, and with root volume and total root number in this prior work. Potri.016G098500 encodes a putative heme-binding protein we found to be associated with the first PC of LRL traits (across root types and timepoints) and that Sun et al. (2019), found associated with LRL (Table S7). Possible reasons for the relatively low level of overlap between these studies include differences between species of poplar, variation in GWAS populations and statistical methods, as well as the use of different rooting assays (Sun *et al*., 2019).

## Conclusion

We performed GWAS to identify regulators of adventitious rooting capacity in 1,148 genotypes from a *P. trichocarpa* clone bank. To facilitate the collection of quantitative measures of adventitious root development, we employed a phenotyping system tailored for our adventitious rooting assay in poplar. The hundreds of gene candidates identified include regulators of cell division and structure, hormone signaling, reactive oxygen species signaling and post-translational modifications as well as many genes of miscellaneous or unknown function. The distinct origins of basal and lateral roots were supported both by our multivariate phenotype analysis and GWAS associations. As root development is a complex and polygenic process, future research will benefit from investigation of interactions between genes such as the candidates identified here, functional studies such as through mutagenesis, and differential rooting responses to environmental treatments.

## Acknowledgements

We thank the National Science Foundation Plant Genome Research Program for support (IOS #1546900, Analysis of genes affecting plant regeneration and transformation in poplar), and members of GREAT TREES Research Cooperative at OSU for its support of the Strauss laboratory.

Support for the Poplar GWAS dataset is provided by the U.S. Department of Energy, Office of Science Biological and Environmental Research (BER) via the Center for Bioenergy Innovation (CBI) under Contract No. DE-PS02-06ER64304. The Poplar GWAS Project used resources of the Oak Ridge Leadership Computing Facility and the Compute and Data Environment for Science at Oak Ridge National Laboratory, which is supported by the Office of Science of the U.S. Department of Energy under Contract No. DE-AC05-00OR22725. We would like to thank the efforts of personnel from the CBI in establishing the GWAS resource used for this study.

This work used the COMET high-performance cluster at the San Diego Supercomputing Center (University of California, San Diego) made available through the Extreme Science and Engineering Discovery Environment (XSEDE), which is supported by National Science Foundation grant number ACI-1548562.

## Author Contributions

Strauss, Li, Jiang, and Muchero designed and directed the overall study, and obtained funding for its execution; Ma, Peremyslova, Magnuson, and Goddard designed and/or executed the phenotypic analyses; Nagle, Yuan, and Damanpreet created, adapted, and executed the machine vision, computation, and data analysis pipelines; Niño de Rivera assisted with inspecting results in IGV. Nagle wrote the manuscript with editing from Strauss, and all others contributed further edits and revisions.

## Data Availability

Raw data and code used for this project is available upon request to the authors. MTMC-SKAT is available on GitHub (https://github.com/naglemi/mtmcskat).

## Notes

### Competing Interest Statement

The authors have declared no competing interest.

