## Supplementary material for "GWAS identifies candidate genes controlling adventitious rooting in *Populus trichocarpa*": Fig. S1; Fig. S2

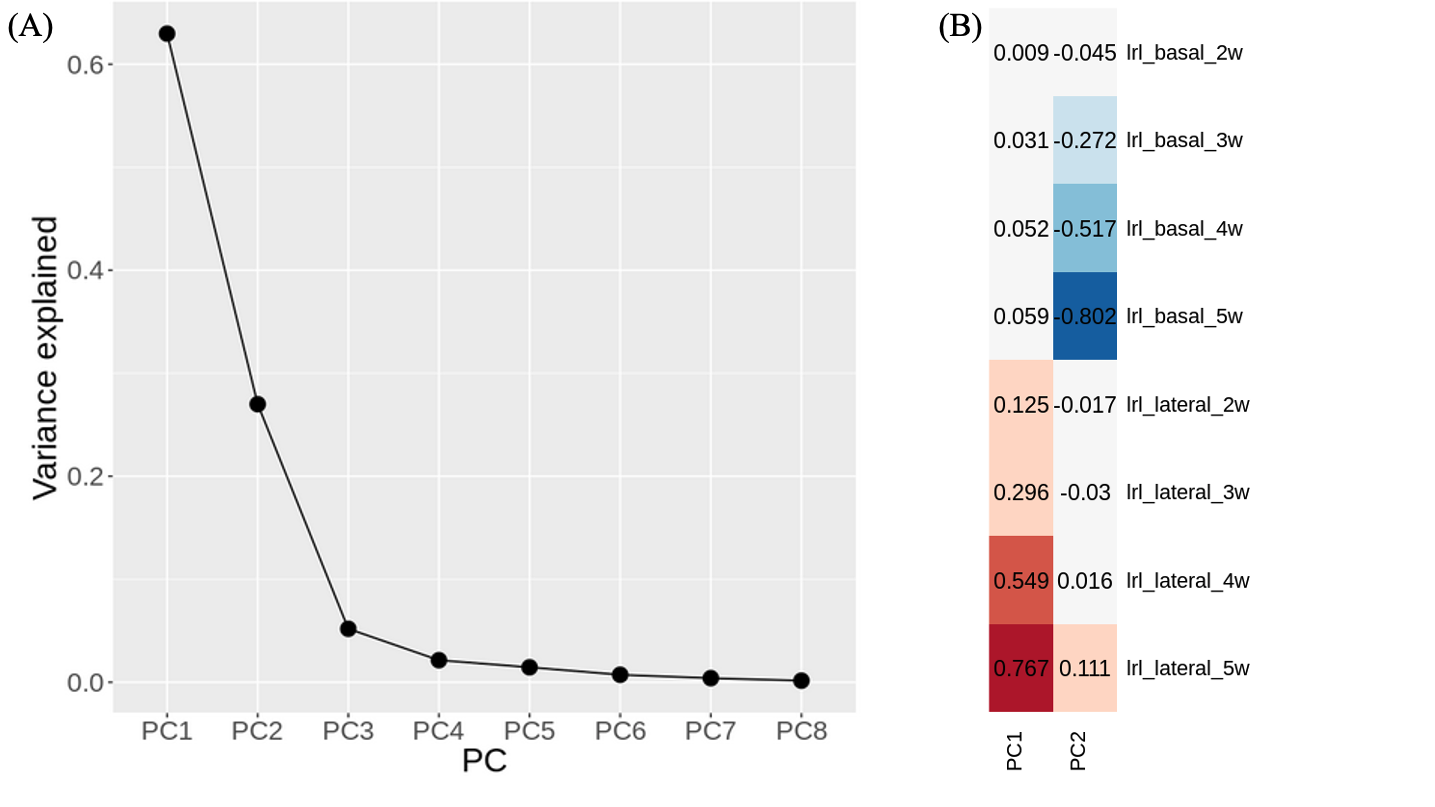

**Supplemental Figure 1.** Results from PCA over all longest root length (LRL) traits, across root type (basal or lateral) and all four timepoints of data collection: (A) Scree plot showing proportion of variance explained by each PC; (B) Loadings for top two PCs.

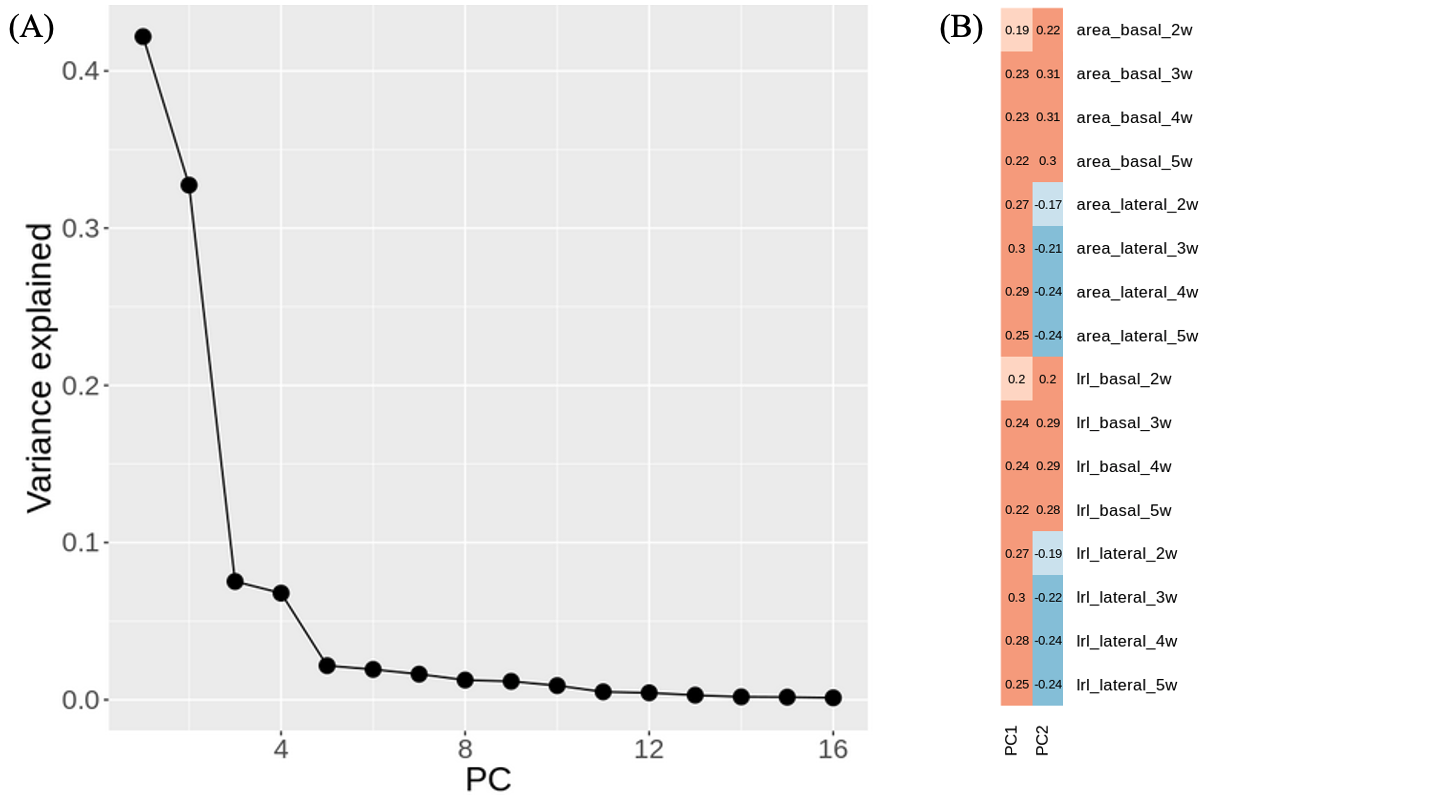

**Supplemental Figure 2.** Results from PCA over all longest root length (LRL) and root area traits, across root type (basal or lateral) and all four timepoints of data collection: (A) Scree plot showing proportion of variance explained by each PC; (B) Loadings for top two PCs.
