## Supplementary material for "GWAS identifies candidate genes controlling adventitious rooting in *Populus trichocarpa*": Table S1; Table S2; Table S3; Table S4

| *Trait category* | *Specific trait* | *Pearson CC of trait and normal dist.* | |
| --- | --- | --- | --- |
|  |  | *Before transf.* | *After transf.* |
| Basal root area | Growth from week 2-5 | 0.667 | 0.994 |
|  | Week 2 | 0.413 | 0.995 |
|  | Week 3 | 0.561 | 0.995 |
|  | Week 4 | 0.612 | 0.996 |
|  | Week 5 | 0.665 | 0.995 |
| Lateral root area | Growth from week 2-5 | 0.893 | 0.998 |
|  | Week 2 | 0.671 | 0.997 |
|  | Week 3 | 0.763 | 0.999 |
|  | Week 4 | 0.846 | 0.996 |
|  | Week 5 | 0.885 | 0.997 |
| Longest basal root | Week 2 | 0.502 | 0.994 |
|  | Week 3 | 0.653 | 0.995 |
|  | Week 4 | 0.696 | 0.992 |
|  | Week 5 | 0.746 | 0.996 |
| Longest lateral root | Week 2 | 0.766 | 0.997 |
|  | Week 3 | 0.857 | 0.998 |
|  | Week 4 | 0.905 | 0.995 |
|  | Week 5 | 0.933 | 0.997 |
| Longest root | Week 2 | 0.782 | 0.997 |
|  | Week 3 | 0.885 | 0.997 |
|  | Week 4 | 0.927 | 0.993 |
|  | Week 5 | 0.954 | 0.996 |
| Total root area | Growth from week 2-5 | 0.907 | 0.998 |
|  | Week 2 | 0.691 | 0.996 |
|  | Week 3 | 0.784 | 0.999 |
|  | Week 4 | 0.852 | 0.998 |
|  | Week 5 | 0.903 | 0.998 |

**Supplementary Table 1.** Raw traits obtained by parsing machine vision outputs into vectors with value of continuous trait for each genotype; all traits were transformed first by excluding zero values, and finally by a Box-Cox transformation.

| *Traits reduced by PCA* | *PC #* | *Transf. type* | *Pearson CC of trait and normal dist.* | |
| --- | --- | --- | --- | --- |
|  |  |  | Before transf. | After transf. |
| Eight root length traits; Longest root length across basal and lateral roots and all four timepoints | PC1 | A | 0.930 | 0.997 |
|  | PC2 | C | 0.790 | 0.993 |
|  | PC3 | B | 0.955 | 0.997 |
|  | PC4 | B | 0.869 | 0.981 |
| Eight root area traits; Root area across basal and lateral roots and all four timepoints | PC1 | A | 0.880 | 0.997 |
|  | PC2 | C | 0.687 | 0.984 |
|  | PC3 | B | 0.905 | 0.995 |
|  | PC4 | B | 0.775 | 0.980 |
| All sixteen root traits; Root area and longest root length across basal and lateral roots and all four timepoints | PC1 | A | 0.879 | 0.999 |
|  | PC2 | B | 0.903 | 0.990 |
|  | PC3 | B | 0.775 | 0.974 |
|  | PC4 | B | 0.942 | 0.996 |

**Supplementary Table 2.** Principal component traits obtained by performing principal component analysis (PCA) to reduce large numbers of raw traits (Table 1) into a smaller number of variables. These traits were processed by a range of transformations, labeled in the “Transf. type” column. Transformation “A” simply involved excluding genotypes which have a zero value for all raw traits, followed by a Box-Cox transformation. Transformation “B” additionally featured removal of outliers prior to Box-Cox, while “C” involved thresholding at an inflection point (elbow) prior to Box-Cox.

| **Trait** | **Heritability** | | **Treatment of trait data before GWAS** | | | |
| --- | --- | --- | --- | --- | --- | --- |
|  | **h^2^_SNP_** | **SE(h^2^_SNP_)** | **Threshold** | **Removal of duplicate values** | **Removal of outliers** | **Transformation** |
| Total root area PC2 | 0.218 | 0.084 | None | Y | N | RB-INV |
| Longest root length PC2 | 0.205 | 0.079 | None | Y | N | RB-INV |
| Total root area (wk. 5) | 0.141 | 0.094 | None | N | N | Box-Cox |
| Total root area growth (wk. 2 - wk. 5) | 0.138 | 0.089 | None | N | N | Box-Cox |
| Total root area PC1 | 0.127 | 0.097 | None | Y | N | RB-INV |
| Root traits overall PC1 | 0.125 | 0.100 | None | Y | N | RB-INV |
| Total root area PC1 | 0.123 | 0.096 | None | Y | N | Box-Cox |
| Longest root length PC1 | 0.120 | 0.094 | None | Y | N | RB-INV |
| Longest root length PC1 | 0.117 | 0.096 | None | Y | N | Box-Cox |
| Root traits overall PC1 | 0.117 | 0.098 | None | Y | N | Box-Cox |
| Basal root area growth (wk. 2 - wk. 5) | 0.114 | 0.135 | None | N | N | Box-Cox |
| Longest lateral root length (wk. 3) | 0.113 | 0.110 | None | N | N | Box-Cox |
| Basal root area (wk. 5) | 0.112 | 0.136 | None | N | N | Box-Cox |
| Root traits overall PC2 | 0.104 | 0.066 | None | Y | N | RB-INV |
| Total root area (wk. 3) | 0.100 | 0.146 | None | N | N | Box-Cox |
| Lateral root area (wk. 3) | 0.075 | 0.135 | None | N | N | Box-Cox |
| Root traits overall PC2 | 0.050 | 0.069 | None | Y | Y | Box-Cox |
| Longest lateral root length (wk. 2) | 0.044 | 0.135 | None | N | N | Box-Cox |
| Longest root length (wk. 5) | 0.037 | 0.120 | None | N | N | Box-Cox |
| Longest basal root length (wk. 5) | 0.021 | 0.108 | None | N | N | Box-Cox |
| Longest root length (wk. 2) | 0.020 | 0.228 | None | N | N | Box-Cox |
| Longest lateral root length (wk. 5) | 0.020 | 0.068 | None | N | N | Box-Cox |
| Basal root area (wk. 3) | 9.37 E-03 | 0.214 | None | N | N | Box-Cox |
| Longest root length PC2 | 5.36 E-03 | 0.078 | 0.708 | Y | Y | Box-Cox |
| Longest root length (wk. 3) | 1.66 E-03 | 0.128 | None | N | N | Box-Cox |
| Total root area PC2 | 2.07 E-06 | 0.279 | 0.364 | Y | Y | Box-Cox |
| Lateral root area (wk. 5) | 2.04 E-06 | 0.062 | None | N | N | Box-Cox |
| Lateral root area growth (wk. 2- wk. 5) | 2.04 E-06 | 0.144 | None | N | N | Box-Cox |
| Longest lateral root length (wk. 4) | 2.04 E-06 | 0.057 | None | N | N | Box-Cox |
| Lateral root area (wk. 4) | 2.04 E-06 | 0.061 | None | N | N | Box-Cox |
| Lateral root area (wk. 2) | 2.04 E-06 | 0.092 | None | N | N | Box-Cox |
| Longest root length (wk. 4) | 2.03 E-06 | 0.078 | None | N | N | Box-Cox |
| Total root area (wk. 4) | 2.03 E-06 | 0.065 | None | N | N | Box-Cox |
| Total root area (wk. 2) | 2.03 E-06 | 0.075 | None | N | N | Box-Cox |
| Longest basal root length (wk. 3) | 1.96 E-06 | 0.235 | None | N | N | Box-Cox |
| Longest basal root length (wk. 4) | 1.96 E-06 | 0.220 | None | N | N | Box-Cox |
| Basal root area (wk. 4) | 1.96 E-06 | 0.411 | None | N | N | Box-Cox |
| Longest basal root length (wk. 2) | 1.95 E-06 | 0.331 | None | N | N | Box-Cox |
| Basal root area (wk. 2) | 1.95 E-06 | 2.447 | None | N | N | Box-Cox |

**Supplementary Table 3.** Narrow-sense heritability (SNP heritability) is shown as computed by GEMMA.

| **Grouping of QTLs by significance and distance** | **Trait** | **Method** | **N genes** | **Positions of QTL peaks relative to nearest transcript** | | | | |
| --- | --- | --- | --- | --- | --- | --- | --- | --- |
|  |  |  |  | **Avg. distance (bp)** | **Median distance (bp)** | **Percent intergenic** | **Percent upstream** | **Percent downstream** |
| All QTLs passing Bonf. | Basal area growth (wk. 2-5) | MTMCSKAT | 6 | 1,648 | 195 | 50% | 33% | 17% |
|  | Longest lateral root (wk. 3) | MTMCSKAT | 13 | 8,464 | 3,423 | 77% | 23% | 54% |
|  | LRL PC2 | MTMCSKAT | 1 | 0 | 0 | 0% | 0% | 0% |
|  | Total root area PC1 | MTMCSKAT | 9 | 1,722 | 323 | 67% | 44% | 22% |
|  | Total root area PC2 | MTMCSKAT | 4 | 691 | 365 | 50% | 50% | 0% |
|  | All root traits PC1 | MTMCSKAT | 1 | 35,938 | 35,940 | 100% | 100% | 0% |
|  | All root traits PC2 | MTMCSKAT | 1 | 21,355 | 21,360 | 100% | 0% | 100% |
|  | Basal root area (wk. 5) | MTMCSKAT | 6 | 1,770 | 560 | 67% | 50% | 17% |
|  | Total root area (wk. 5) | MTMCSKAT | 1 | 9,499 | 9,499 | 100% | 0% | 100% |
| All QTLs passing Bonf. within 5kb of gene | Basal area growth (wk. 2-5) | MTMCSKAT | 5 | 485 | 0 | 40% | 20% | 20% |
|  | Longest lateral root (wk. 3) | MTMCSKAT | 9 | 1,805 | 1,181 | 67% | 33% | 33% |
|  | LRL PC2 | MTMCSKAT | 1 | 0 | 0 | 0% | 0% | 0% |
|  | Total root area PC1 | MTMCSKAT | 8 | 628 | 188 | 63% | 38% | 25% |
|  | Total root area PC2 | MTMCSKAT | 4 | 691 | 365 | 50% | 50% | 0% |
|  | Basal root area (wk. 5) | MTMCSKAT | 5 | 631 | 390 | 60% | 40% | 20% |
| All QTLs passing FDR (alpha = 0.10) and/or Bonf. | Basal area growth (wk. 2-5) | MTMCSKAT | 28 | 13,475 | 3,206 | 68% | 43% | 25% |
|  | Longest lateral root (wk. 3) | GEMMA | 2 | 8,837 | 8,836 | 100% | 50% | 50% |
|  | Longest lateral root (wk. 3) | MTMCSKAT | 65 | 6,078 | 1,936 | 77% | 40% | 37% |
|  | LRL PC1 | MTMCSKAT | 10 | 3,254 | 2,996 | 80% | 40% | 40% |
|  | LRL PC2 | MTMCSKAT | 5 | 16,432 | 5,659 | 80% | 40% | 40% |
|  | Total root area PC1 | MTMCSKAT | 29 | 5,837 | 1,437 | 69% | 34% | 34% |
|  | Total root area PC2 | MTMCSKAT | 26 | 10,690 | 2,470 | 77% | 46% | 31% |
|  | All root traits PC1 | MTMCSKAT | 29 | 12,084 | 1,942 | 66% | 34% | 31% |
|  | All root traits PC2 | MTMCSKAT | 6 | 4,129 | 617 | 83% | 0% | 83% |
|  | Basal root area (wk. 5) | MTMCSKAT | 21 | 10,773 | 6,570 | 86% | 57% | 29% |
|  | Total root area (wk. 5) | MTMCSKAT | 2 | 5,938 | 5,938 | 100% | 50% | 50% |
| All QTLs passing FDR (alpha = 0.10) and/or Bonf. within 5kb of gene | Basal area growth (wk. 2-5) | MTMCSKAT | 17 | 1,050 | 0 | 47% | 35% | 12% |
|  | Longest lateral root (wk. 3) | MTMCSKAT | 48 | 1,350 | 617 | 69% | 40% | 29% |
|  | LRL PC1 | MTMCSKAT | 7 | 1,565 | 608 | 71% | 43% | 29% |
|  | LRL PC2 | MTMCSKAT | 2 | 2,484 | 2,484 | 50% | 50% | 0% |
|  | Total root area PC1 | MTMCSKAT | 23 | 1,267 | 575 | 61% | 39% | 22% |
|  | Total root area PC2 | MTMCSKAT | 16 | 1,076 | 411 | 63% | 38% | 25% |
|  | All root traits PC1 | MTMCSKAT | 18 | 900 | 0 | 44% | 28% | 17% |
|  | All root traits PC2 | MTMCSKAT | 5 | 684 | 396 | 80% | 0% | 80% |
|  | Basal root area (wk. 5) | MTMCSKAT | 10 | 938 | 560 | 70% | 50% | 20% |
|  | Total root area (wk. 5) | MTMCSKAT | 1 | 2,377 | 2,377 | 100% | 100% | 0% |
| All QTLs passing ART-Bonf. | Basal area growth (wk. 2-5) | GMMAT | 4 | 4,127 | 3,118 | 100% | 50% | 50% |
|  | Total area growth (wk. 2-5) | GMMAT | 3 | 1,444 | 840 | 100% | 67% | 33% |
|  | Total root area PC2 | GMMAT | 1 | 1,695 | 1,695 | 100% | 0% | 100% |
|  | Basal area growth (wk. 2-5) | GEMMA | 11 | 3,583 | 1,696 | 91% | 36% | 55% |
|  | Total root area growth (wk. 2-5) | GEMMA | 17 | 2,445 | 946 | 82% | 29% | 53% |
|  | Longest lateral root (wk. 3) | GEMMA | 11 | 3,163 | 1,932 | 91% | 27% | 64% |
|  | LRL PC1 | GEMMA | 13 | 2,800 | 2,740 | 100% | 54% | 46% |
|  | LRL PC2 | GEMMA | 15 | 3,490 | 2,893 | 87% | 53% | 33% |
|  | Total root area PC1 | GEMMA | 16 | 2,335 | 1,712 | 88% | 44% | 44% |
|  | Total root area PC2 | GEMMA | 17 | 2,104 | 528 | 82% | 29% | 53% |
|  | All root traits PC1 | GEMMA | 4 | 3,105 | 3,068 | 100% | 75% | 25% |
|  | All root traits PC2 | GEMMA | 11 | 4,665 | 2,130 | 100% | 36% | 64% |
|  | Basal root area (wk. 5) | GEMMA | 12 | 3,658 | 2,210 | 83% | 42% | 42% |
|  | Total root area (wk. 5) | GEMMA | 7 | 1,791 | 777 | 86% | 43% | 43% |
| All QTLs passing ART-Bonf. within 5kb of gene | Basal area growth (wk. 2-5) | GMMAT | 2 | 987 | 987 | 100% | 50% | 50% |
|  | Total area growth (wk. 2-5) | GMMAT | 3 | 1,444 | 840 | 100% | 67% | 33% |
|  | Total root area PC2 | GMMAT | 1 | 1,695 | 1,695 | 100% | 0% | 100% |
|  | Basal area growth (wk. 2-5) | GEMMA | 7 | 1,235 | 1,109 | 86% | 29% | 57% |
|  | Total root area growth (wk. 2-5) | GEMMA | 14 | 1,258 | 898 | 79% | 29% | 50% |
|  | Longest lateral root (wk. 3) | GEMMA | 8 | 1,575 | 1,578 | 88% | 38% | 50% |
|  | LRL PC1 | GEMMA | 12 | 2,117 | 2,293 | 100% | 58% | 42% |
|  | LRL PC2 | GEMMA | 11 | 2,051 | 1,892 | 82% | 55% | 27% |
|  | Total root area PC1 | GEMMA | 15 | 1,823 | 1,579 | 87% | 40% | 47% |
|  | Total root area PC2 | GEMMA | 15 | 1,081 | 505 | 80% | 33% | 47% |
|  | All root traits PC1 | GEMMA | 3 | 2,085 | 2,737 | 100% | 67% | 33% |
|  | All root traits PC2 | GEMMA | 8 | 1,684 | 1,536 | 100% | 25% | 75% |
|  | Basal root area (wk. 5) | GEMMA | 7 | 1,004 | 1,041 | 71% | 29% | 43% |
|  | Total root area (wk. 5) | GEMMA | 6 | 1,101 | 586 | 83% | 33% | 50% |

**Supplementary Table 4.** Tallies and statistics of QTL peaks found significant across methods, traits and significance thresholds
